# Insight into the emerging insect to human pathogen *Photorhabdus* revealing geographic differences in immune cell tropism

**DOI:** 10.1101/2023.08.22.554382

**Authors:** Max L. Addison, Alexia Hapeshi, Elena Carter, Zi Xin Wong, John E. Connally, Nicholas R. Waterfield

## Abstract

*Photorhabdus asymbiotica* is a species of the insect pathogenic *Photorhabdus* genus that has been isolated as an etiological agent in human infections. Since then, multiple isolates have been identified worldwide, however actual clinical infections have so far only been identified in North America, Australia and Nepal. Previous research on the clinical isolates had shown that the strains differed in their behaviour when infecting cultured human cells, based on the geographical site of isolation. In this study we investigate the differences between the pathogenic activities of *P. asymbiotica* isolates from different geographic locations. We present findings from the clinical isolates from Australia (Kingscliff) and North America (ATCC43949), and soil borne nematode isolates from Thailand (PB68) and Northern Europe non-clinical *P. asymbiotica* genospecies HIT and JUN. We also show the first findings from a new clinical isolate of *P. luminescens* (Texas), the first non-*asymbiotica* species to cause a human infection, confirming its ability to infect and survive inside human immune cells. Infection assays were done using both cultured cell lines (THP-1, CHO and HEK cells) and also primary immune cells, Peripheral Blood Mononuclear cells (PBMCs) isolated from human blood. Here for the first time, we show how *P. asymbiotica*, selectively infects certain immune cells while avoiding others, and that infectivity varies depending on growth temperature. We also show that the infected immune cells vary depending on the geographical location a strain is isolated from, and that the European HIT and JUN strains lack the ability to survive within mammalian cells in tissue culture.

## Introduction

Members of the genus *Photorhabdus* are entomopathogenic bacteria, which form an obligate symbiosis with the insect parasitic nematode, *Heterorhabditis*. The free-living stage of the nematodes, infective juveniles, are found in the soil carrying *Photorhabdus* in their gut, where they search for insect larvae. Once found they will burrow into the insect before regurgitating the *Photorhabdus* which essentially acts as a “bioweapon”. The bacteria release a large range of toxins and enzymes which kill the insect allowing them to replicate. The nematode then begins hermaphrodite replication cycles, using the bacterial biomass as a food source. The speed and efficiency of the nematode-*Photorhabdus* partnership in killing insect larvae, generally under 48 hours (Frost et al, 1997), has made it an effective and widely used bio-pesticide.

While most species of *Photorhabdus* are only able to infect insects, being unable to grow above 34 °C, one species has been reported to cause human infections, *Photorhabdus asymbiotica*. Like their insect restricted relatives, the bacteria are carried by symbiont *Heterorhabditis* nematodes. Of all reported cases of *Photorhabdus* infections only one case did not involve a member of the *P. asymbiotica* species. In this case an isolate of a novel strain of *P. luminescens* was found to have infected a neonate in Texas, that was abandoned just hours after birth on a patch of soil. This strain is currently poorly characterised however, so it is unknown if it would be able to infect an adult, or merely took advantage of the neonate’s weakened immune system. It is perhaps telling however that this strain, unlike the majority of other *P. luminescens* isolates, can grow at 37 °C. Infections of *Photorhabdus*, known as Photorhabdosis, typically start with a large lesion generally on an extremity believed to be initial site of infection. If untreated secondary lesions appear on other parts of the body and in some cases, bacteria were found in the respiratory system and the heart causing endocarditis, confirming the ability of *Photorhabdus* to disseminate throughout body. It has been hypothesised they may achieve this by hijacking immune cells and essentially “hitchhiking” via the lymphatic system. This facultative intracellular infection behaviour is seen in the phylogenetically closely related *Yersinia pestis*, which are carried inside phagocytes around the body (St John *et al*, 2014). It is unknown how *Photorhabdus* enters the body of the human host, however the most likely hypothesis is that an infective juvenile accidently burrows into the dermal layer of the human skin. While the *Photorhabdus* can establish an infection, the nematode would not likely be able to survive core body temperature and would need to remain near the cooler surface of the host. Some related nematodes such as *Strongyloides stercoralis* have been shown to be able to burrow through human skin (Gang *et al*, 2020). Nevertheless, a *Heterorhabditis* nematode has never been isolated from one of the human infections, although no one has purposely looked for them to date.

**Figure 1.**
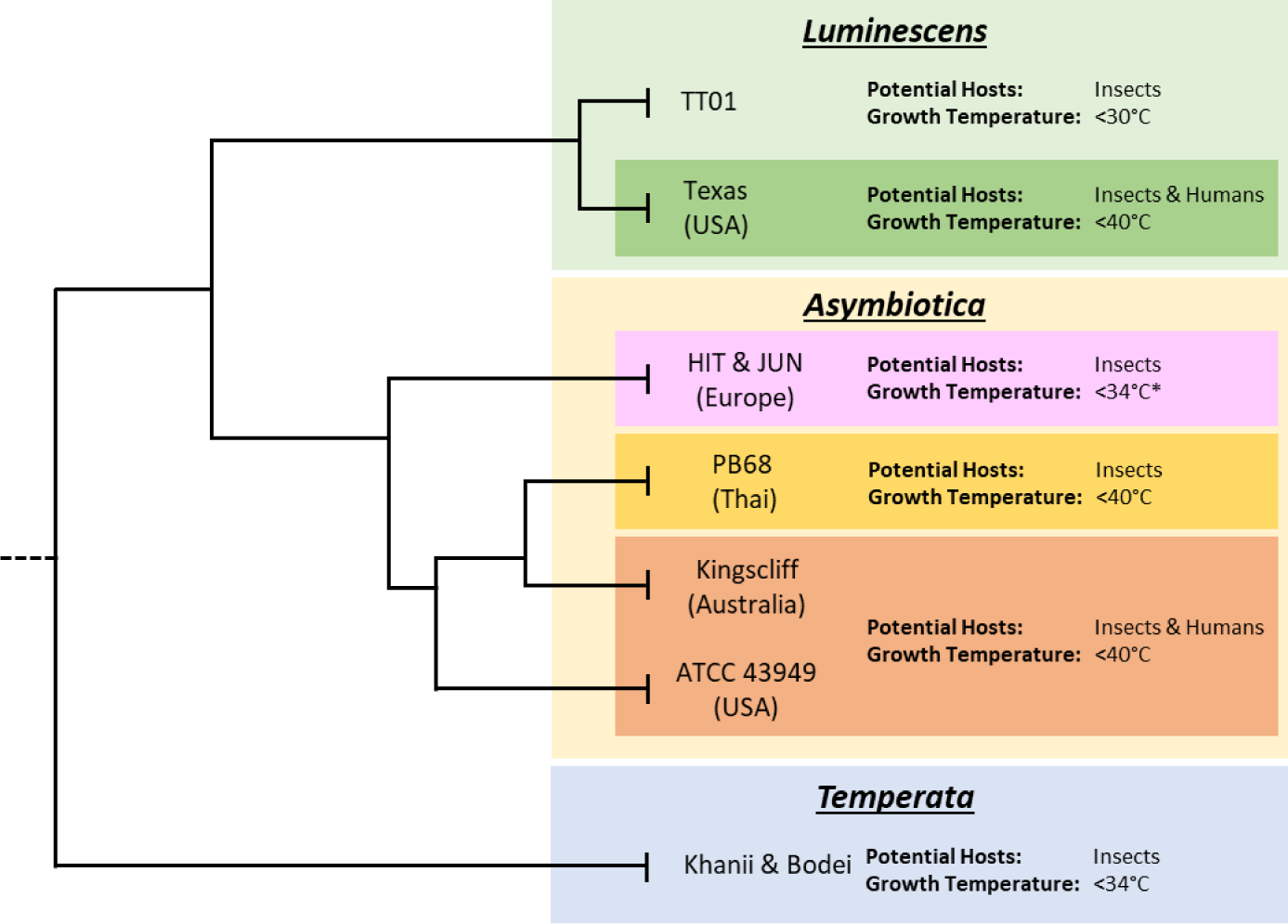
Subclades of the various *Photorhabdus* species. Tree lengths are not drawn to scale and for illustrative purposes only. Displayed are the approximate thermotolerances and known potential hosts of archetypal strains of *Photorhabdus* within each species. Data on thermotolerances is from (Mulley et al, 2015) and our own observations from working with our strain collection. While published data on the HIT and JUN strains showed inability to grow at 37°C, during this project it was found that given extended growth times, of 48 hours compared to 24 hours, they would regularly reach higher ODs and eventually reach stationary phase.

Interestingly so far incidences of human infection have been geographically isolated, only being reported in eastern Australia, continental eastern North America and a single case contracted by a traveller in Nepal (Farmer *et al*, 1989; Pujol & Bliska, 2005). In these cases, many of the infections were clustered in geographically constrained areas of those countries, for example, Texas in North America and Victoria in Australia. Both these areas have sandy soils which the *Heterorhabditis* nematodes seem to prefer, and infections seem to coincide with warm, wet weather, which may stimulate nematode activity or perhaps insect host availability. While the number of recognised human infections is low, it should be noted that until as recently as 2017, *Photorhabdus* was not included in the standard databases used by many medical professionals to identify infections. This may have led to it being commonly mistakenly identified as a different species of bacteria. When tested in a lab setting using a VITEK 2 Gram-negative identification card it indeed did lead to misidentification, in this case as *Pseudomonas fluorescens* (Weissfeld *et al*, 2005). The most accurate way to identify *Photorhabdus* infections is through the use of 16S ribosomal DNA gene or *recA* sequencing. However, this is time consuming, typically taking 48-72 hours (Boyles and Wasserman, 2015). A simple alternative is by visible inspection for bioluminescence by dark adapted eyes as *Photorhabdus* is the only known terrestrial bioluminescent bacterium. These factors, along with the ease that most *Photorhabdus* infections can be treated, using a standard antibiotic course, means that many *Photorhabdus* infections are likely to have gone underdiagnosed. This may be the reason why we have only had reports of infections in the USA and Australia, as other places where *P. asymbiotica* has been found, such as rural Thailand, may not have the resources, infrastructure or requisite knowledge to properly identify infections. In reality infection clusters may actually only reflect the presence of clinicians who are aware of this pathogen.

*P. asymbiotica* strains have all been reported to grow at human body temperature, with some strains being able to grow at up to 42°C. However, while it has been a long time since the first recorded case of a *Photorhabdus* human infection, 1977 (Farmer *et al*, 1989), there has been little study on this emerging human pathogenic species of *Photorhabdus.* Thus, it is unknown if temperature tolerance is all that separates the human infective strains from the non-human infective strains. Some differences can be found in the genomes between the *P. asymbiotica* and the non-human infective *P. luminescens* strain TT01. For example, *P. asymbiotica* encodes a more limited range of insect toxicity genes, resulting in a smaller genome, approximately 600,000 base pairs [bp] less than the *P. luminescens* type strain TT01. Notably *P. asymbiotica* lacks a homologue of the type 3 secretion system (T3SS) effector *lopT* gene possessed by all *P. luminescens* so far examined. The T3SS LopT has been shown to prevent phagocytosis (Brugirard-Ricaud *et al*, 2005). In the same locus as *lopT*, *P. asymbiotica* instead carries a homologue of the *exoU* gene from *Pseudomonas aeruginosa*. The ExoU toxin is a phospholipase associated with acute lung injury (Pankhaniya *et al*, 2004).

The only research published on the interaction between *P. asymbiotica* and tissue culture cells confirmed that it is capable of invading human immune cells and resists killing by humoral factors such as the complement system (Costa *et al*, 2009). This research did bring up an interesting observation however, that the American and Australian strains of *P. asymbiotica* differed in their ability to invade human cells, despite both being human pathogenic strains. This raises the question of whether geographically distinct strains of *Photorhabdus* differ in their strategies for evading/resisting the immune system response. So far, all the research on *P. asymbiotica* has only been done using single isolates of the confirmed human infective American and Australian strains. It should also be noted that previous research into *P. asymbiotica* was performed using *Photorhabdus* cultures that had been grown at 28°C prior to cell challenge experiments. In previous studies *Photorhabdus* has been shown to only express certain genes when grown at human body temp of 37 °C compared to the more insect-relevant body temp of 28 °C (Hapeshi *et al*, 2020). This presents the distinct possibility that the initial growth temperature prior to infection may influence the bacterial behaviour and its ability to establish an infection.

To get a clearer understanding of how geographically distinct strains of *Photorhabdus asymbiotica* differ in their strategies for evading/resisting the immune system and how this behaviour is affected by growth temperature, five geographically distant *P. asymbiotica* strains were studied alongside the newly identified human infective *P. luminescens* strain (**Fig. 2**). The strains studied were the North American *P. asymbiotica* subsp *P. asymbiotica* strain ATCC43949, the Australian *P. asymbiotica* subsp. *Australis* strain (Kingscliff), the closely related Thailand isolate *P. asymbiotica* subsp. *Australis* strain (PB68), the two Northern European *P. asymbiotica* subsp, designated HIT and JUN, and finally the recent clinical isolate of *P. luminescens* (Texas). Kingscliff, ATCC43949 and Texas are clinical isolates from confirmed human infections, while the other *P. asymbiotica* genospecies strains were isolated from soil dwelling nematodes, and thus it is unknown if they are capable of causing clinical infections. The lab passaged *P. luminescens* TT01-DJC strain was also included as a non-human infective negative control. To understand how the behaviour of these strains compared, they were tested for their ability to survive or avoid phagocytosis in the human monocyte derived macrophage model cell line, THP-1. We also examined how they interacted with a more natural and complete model of the human immune system. In order to do this, we studied the interactions of these strains with human derived PBMCs [Peripheral blood mononuclear cells] taken from healthy human volunteers (Singapore).

**Figure 2.**
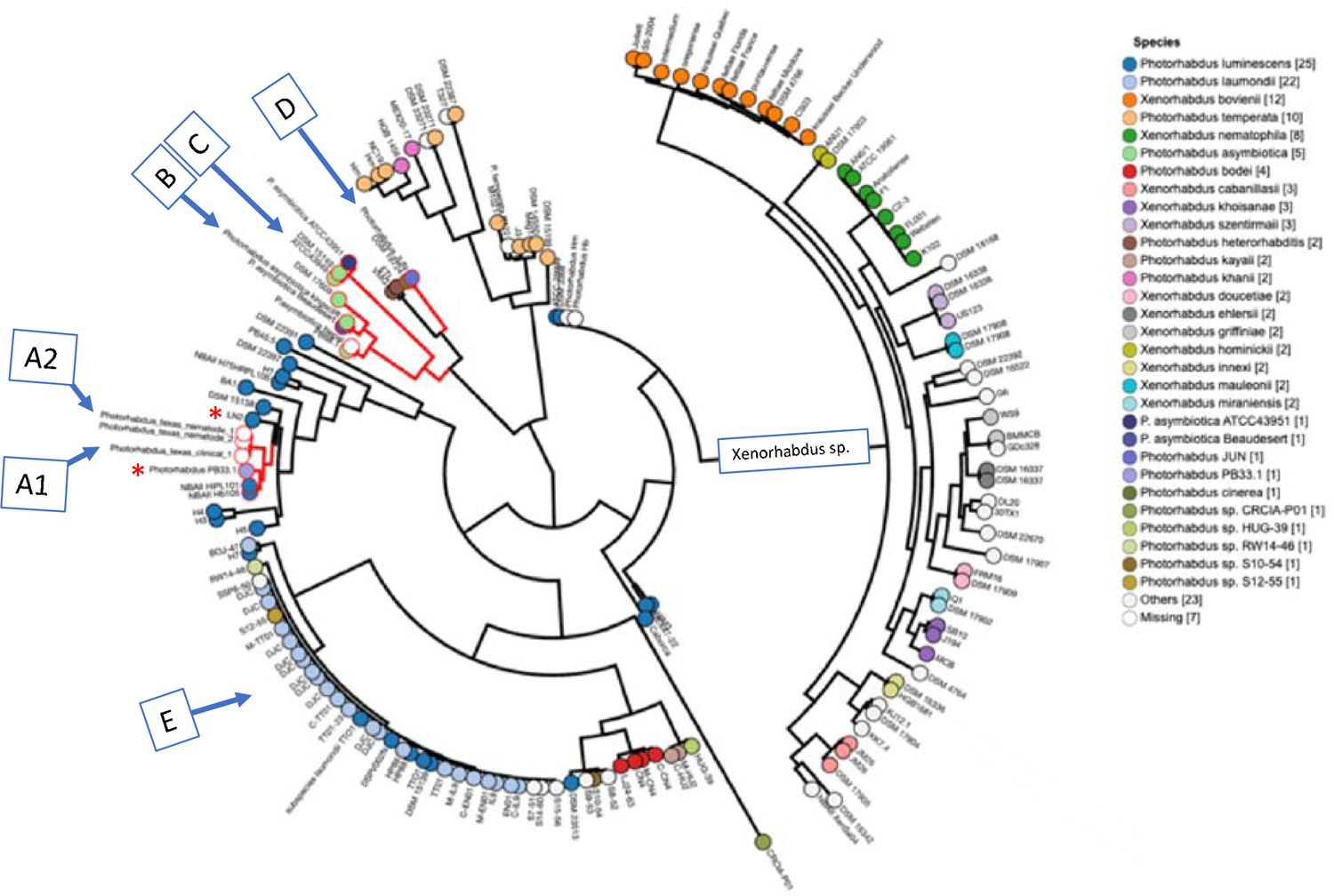
Subclades within the *Photorhabdus* species. Tree lengths are not drawn to scale and for illustrative purposes only. A circular dendrogram showing the phylogeny of *Photorhabdus* and related *Xenorhabdus* sequenced strains available in the Enterobase *Photorhabdus* database (unpublished). Clade branches in red contain strains relevant to this study. [A1] and [A2] are clinical and soil nematode isolates of *P. luminescens* “Texas”; [B] indicates *P. asymbiotica* Kingscliff Australian isolate; [C] indicates *P. asymbiotica* ATCC43949 USA isolate’, [D] indicates *P. asymbiotica* genospecies JUN isolate; [E] indicates *P. luminescens* TT01 derived lab strain DJC. Note, like the “Texas” strains, the of *P. luminescens* PB33.1 and LN2 strains (*) can also grow well at 37 °C.

Here for the first time, we reveal how different strains of *P. asymbiotica*, selectively associate with certain immune cell types while avoiding others. We also show that the range of immune cells that become infected varies depending on the geographical location that the strain was isolated from. This study also confirmed the hypothesis that the Northern European genospecies strains of *P. asymbiotica*, HIT and JUN, lack the ability to survive within mammalian cells, indicating that unlike the previous assumption, not all members of *P. asymbiotica* genospecies clade are capable of infecting mammals. We also show evidence that the clinical strain of *P. luminescens* varies greatly in its behaviour compared to that of the non-human infective *P. luminescens*, but still acts differently to the *P. asymbiotica*.

**Fig. 3.**
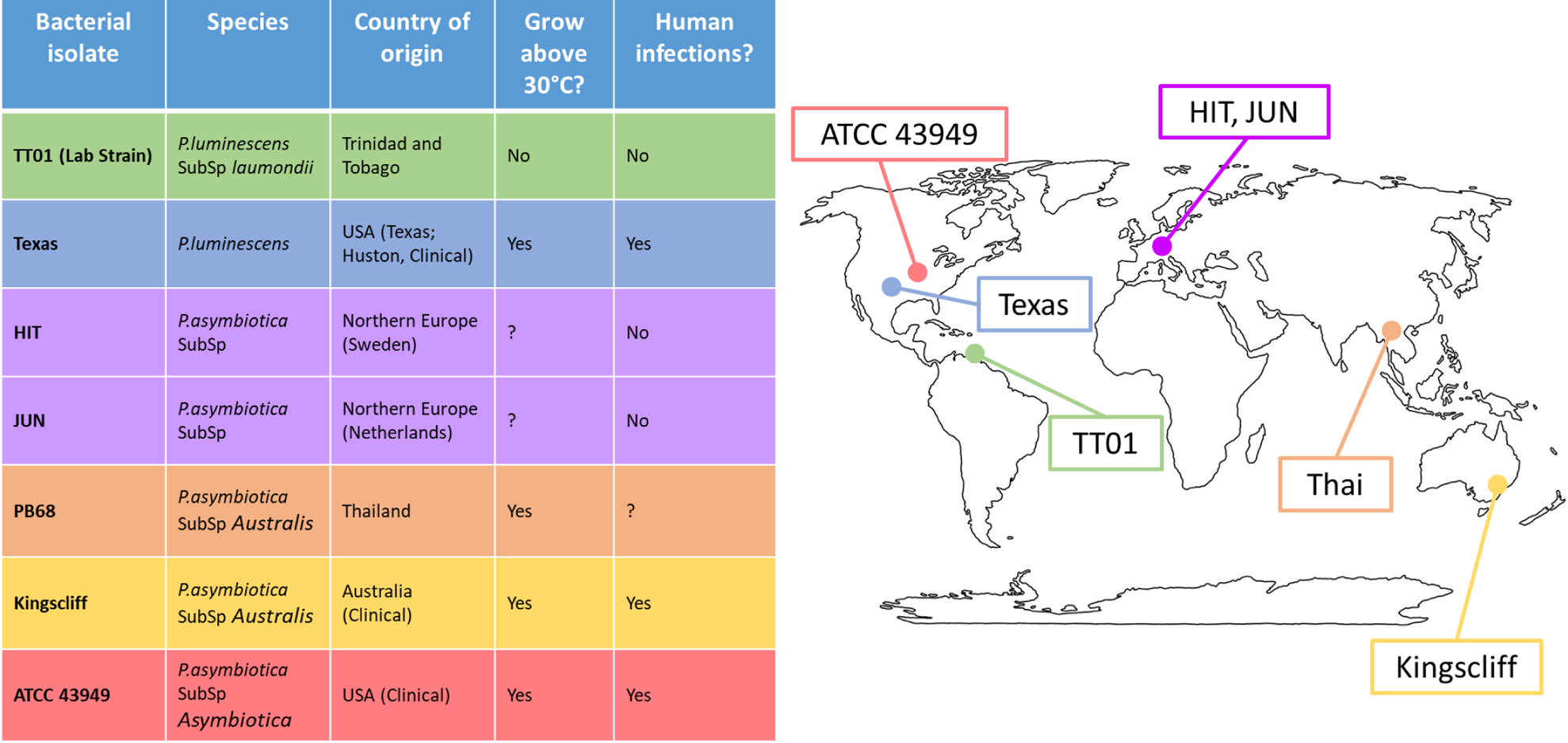
Global distribution of *Photorhabdus* strains used in this study. Table shows The *Photorhabdus* isolates used for the experiments in this study. (Clinical) indicates the isolate used was isolated directly from a human infection, otherwise the isolate was isolated from soil dwelling nematodes in the region. A question mark indicates that not enough research has been done previously to confirm or deny the attribute.

## Results

### Both the growth temperature and the specific strain of *P. asymbiotica* greatly influences their ability to invade cultured mammalian cells

We studied the effect of growth temperature to afford a basic understanding of the behavioural host cell tropism between different *P. asymbiotica* strains. The five strains investigated (see above) were tested for their ability to invade and/or survive inside human THP-1 cells, which had previously been differentiated into a macrophage like cell line, using gentamycin protection assays. In brief, gentamycin protection assays provide information on the ability of bacteria to invade and survive within cells. In this case the bacteria were incubated with the eukaryotic cells at a multiplicity of infection (MOI) of 1:50 (cell:bacteria) for 2 hours, after which any external bacteria still exposed to the surrounding media were killed using gentamycin (which cannot enter the eukaryotic cell). Subsequently the gentamycin was removed by washing, and the eukaryotic cells lysed, releasing any intracellular bacteria. These were enumerated by colony counts on agar after 24h. Gentamycin protection assays provide an indicator of the ability of bacteria to be internalised and survive within a host cell, as any bacteria remaining exposed to the extracellular milieu are killed by the antibiotic. Prior to exposure to the differentiated THP-1 cells, the bacteria were grown either at 28 °C or 37 °C to mid log phase (approximate OD_600_ = 0.4-0.6. In addition to the THP-1 cell gentamycin protection assays, we also examined interactions with the human kidney like cell line HEK-293T and the hamster ovarian line CHO. As HEK 293T and CHO cells are not professional phagocytes, these assays provided information on the bacteria’s ability to actively invade cells, as opposed to simply being able to escape killing when phagocytosed.

An important observation from these experiments is that prior growth temperature of the bacteria strongly effected the ability of certain strains of *P. asymbiotica* to invade these representative mammalian cells (**Fig.4**).

**Fig. 4.**
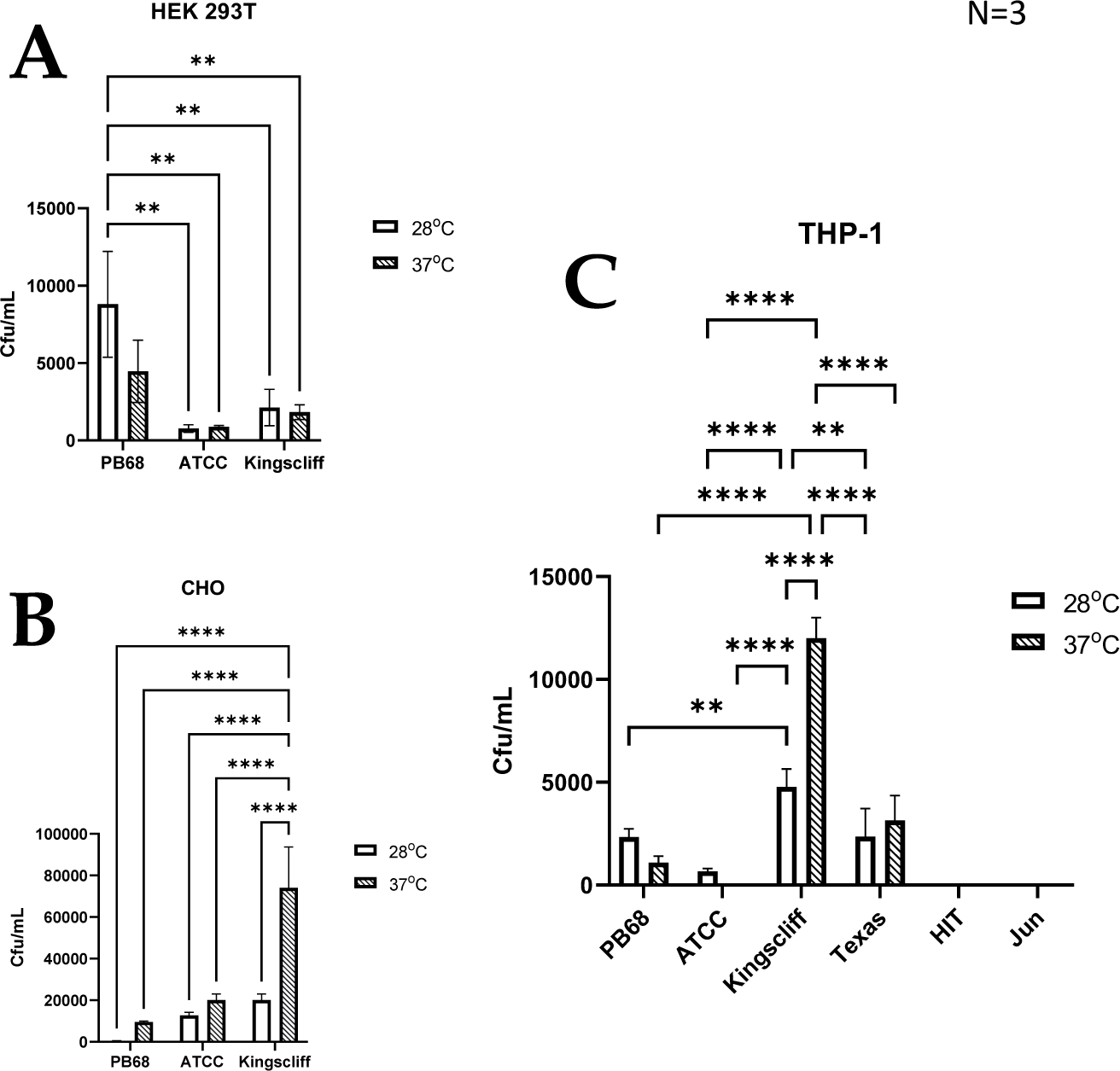
Invasion and survival of *P. asymbiotica* species inside mammalian cells. Gentamycin based invasion assays were performed on several mammalian cell lines using the different bacterial isolates. Bacteria were grown at either 28 °C or 37 °C prior to assay. During the assay the bacteria were allowed to interact with the mammalian cell lines for 2 hours, after which gentamycin was added to kill any extracellular bacteria. The mammalian cells were subsequently lysed, and surviving (presumably intracellular) bacterial numbers assessed using colony forming count plating assays. The infection assays were done with; **A –** HEK 293T, **B- CHO** cells, and **C –** THP-1 cells that had been differentiated into macrophage like cells using PMA, prior to infection. (2way Anova, ** = < 0.01, **** = < 0.0001, n = 3 biological replicates).

An analysis of the cell type tropisms and “invasion” rates of the different *P. asymbiotica* strains supported previous published data on the interaction of *P. asymbiotica* with THP-1 cells. For example, the Australian Kingscliff strain had significantly higher invasion rates than the American ATCC 43949 strain, or indeed all the other *P. asymbiotica* strains tested, regardless of the prior growth temperature (**Fig.4 A**). The increased ability to invade mammalian cells was consistent with the HEK 293T cell assays, however this was only the case if the bacteria were grown at 37 °C prior to the challenge. Interestingly, in CHO cells, Kingscliff exhibited the same invasion rates as ATCC 43949. Having said this, both strains showed significantly lower invasion rates than the Thai PB68 strain which, in the other cells types, mirrored the behaviour of ATCC 43949. The European strains HIT and JUN demonstrated no ability to invade or survive in the host cells, which is perhaps not surprising given their inability to easily grow and replicate above 34 °C.

Interestingly the *P. luminescens* Texas strain was also able to survive within the THP-1 cells, supporting the hypothesis that it is indeed an emerging human pathogen. Its internalisation and survival rates were similar to that of PB68, being slightly higher though not significantly so. While the data is not shown here the lab strain of *P. luminescens* TT01-DJC, was not able to survive within the phagocytes. Though this is likely due to it being unable to replicate at the required THP-1 incubation temperatures.

Despite being a clinical isolate, temperature did not have a significant effect on the ability of the ATCC43949 strain to invade cells. Conversely, growing the Kingscliff strain at 37 °C prior to exposure to host cells, significantly increased the ability for it to invade both HEK and THP-1 cells. Interestingly while the invasion propensity of Kingscliff previously grown at 28 °C into the THP-1 cells was higher than the other *P. asymbiotica* strains, when exposed to HEK cells the invasion rate of 28 °C grown cells was similar to that of the others strains, only becoming significantly higher when grown at 37 °C. Strain PB68 showed an opposite trend in comparison to that of Kingscliff, exhibiting a slight increase in invasion propensity when grown at 28 °C, in comparison to when it was grown at 37 °C. Nevertheless, these differences in trends did not prove significant in these experiments.

### Temperature dependant internalisation and /or attachment of different strains of *P. asymbiotica* with THP-1 derived macrophages

While the invasion assay findings described above revealed specific temperature dependant tropisms of the different *Photorhabdus* strains they do not give a truly representative picture of what likely happens in a real infection. These experiments only tell us if the bacteria are being internalised and surviving over a short time period in highly artificial tissue culture conditions. Therefore, we decided to use microscopy to investigate the interaction of the strains with these phagocytic THP-1 derived macrophage cells in more detail. To this end the gentamycin protection assays were repeated as before, but instead of lysing the phagocytes after the incubation time, the samples were imaged using an inverted fluorescent microscope. For these experiments we used GFP labelled variants of the different *P. asymbiotic*a strains. While this was possible for Kingscliff and PB68, we were not able to transform the European strains with the marker plasmid. We therefore used the non-fluorescent wild-type strains for assays with these strains and relied on phase contrast imaging to reveal their location. From the previous infection assays, it may have been expected that the European HIT and JUN strains might either simply attach to the surface of the THP-1 cells or not interact at all. The latter hypothesis was indeed the case for the JUN, strain, where at either temperature the bacteria could not be seen to associate with the THP-1 cells. This may suggest that they are actively avoiding the THP-1 cells (**Fig.5**). Interestingly however, the HIT strain was seen to attach to the THP-1 cells, when grown at either temperature, with possible internalisation at 37°C, although without a fluorescent label we cannot be confident about that observation (**Fig.5**).

**Fig. 5.**
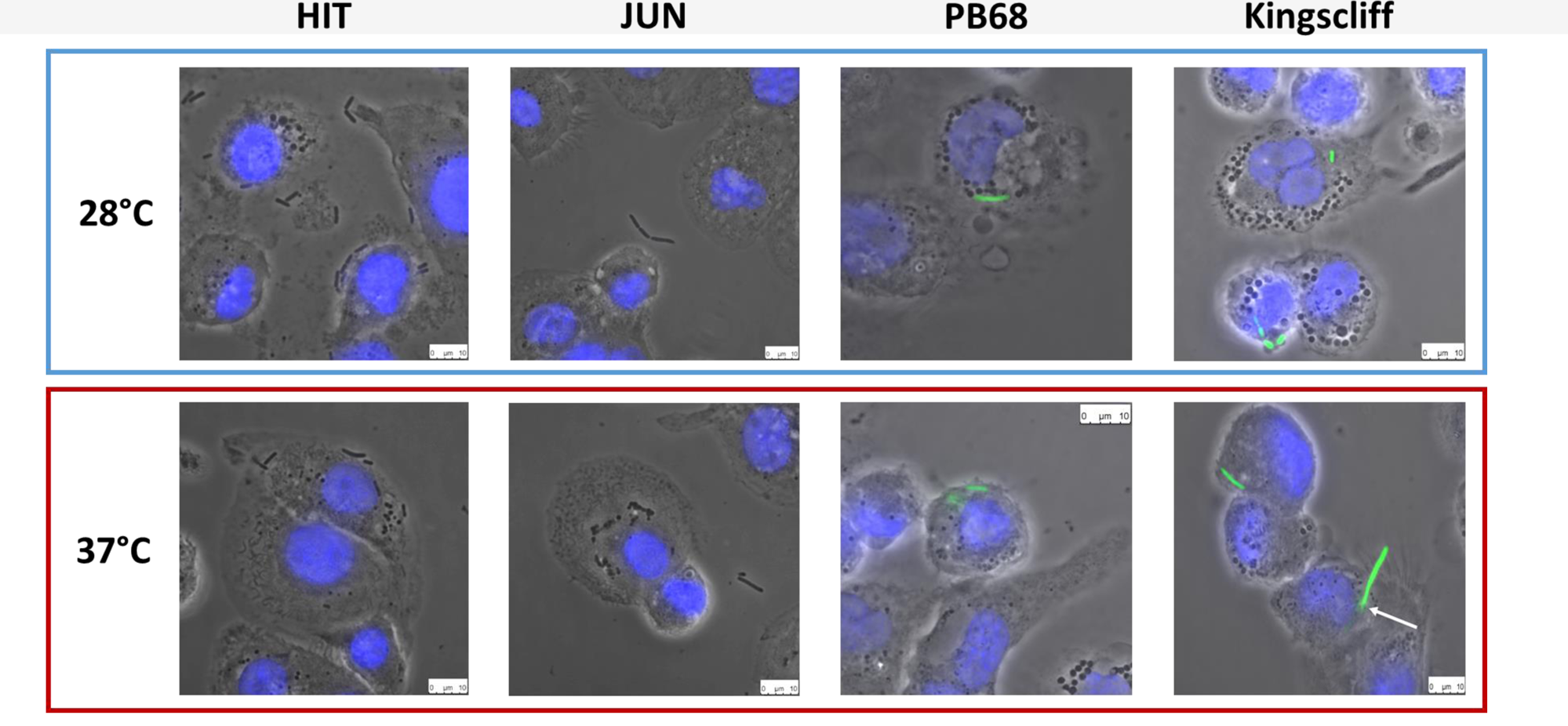
Phase contrast images of *Photorhabdus asymbiotica* subsp trains; HIT, and JUN and fluorescent imaging of Kingscliff and PB68 with THP-1 cells. Before the experiment the THP-1 cells were seeded into 24 well plates on glass coverslips, and differentiated into macrophage like cells using PMA. Bacteria were grown O/N at either 28°C or 37°C prior to infection and allowed to infect for 2 hours, at 37°C, at a MOI of 1:50. After infection the THP-1 cells were washed to remove non-internalised/attached bacteria prior to imaging. THP-1 cell nuclei were stained with DAPI (blue) while the Kingscliff and PB68 strains were constitutively expressing GFP (green).

Conversely, microscopy studies of the infection assays for both Kingscliff and PB68 clearly demonstrated that they were being internalised in the THP-1 cells when previously cultured at either temperature (**Fig.5**). A further interesting observation was that if Kingscliff was grown at 28°C, on occasion long bacterial filaments could be seen apparently crossing the THP-1 cell membrane, indicating that it is either invading or emerging from the cell (**Fig.5**). This leads us to suggest it is using an active mechanism for invasion rather than simple phagocytosis/escape by the THP-1 cell. It was not clear from these assays if the observed bacterial filaments represented chains of single cells or single long multinucleate “hyphae”.

While there are examples of bacteria that can escape the phagosome after engulfment, another strategy often seen in various pathogens demonstrate that they have adaptations which allow them to remain inside the vesicles but prevent phagolysosome maturation and so avoid their destruction. In previous unpublished work with an Austrian isolate of *P. asymbiotica*, researchers commented that transmission electron microscopy was used to confirm that in the insect professional phagocytes (haemocytes), this strain could be observed within phagosomes for at least ∼2 hours post infection. However, the light microscopy methods we used in this study could not discern if the bacteria were enclosed in phagosome vesicles or free in the cytoplasm.

### Internalisation of *P. asymbiotica* requires actin rearrangement

As we confirmed that certain *P. asymbiotica* species are capable of entering a range of cell types, we wanted to better understand the processes involved in cell entry. Many bacteria facilitate uptake and entry into host cells by manipulating host cell actin rearrangements. In non-phagocytic host cells this involves using either so called “zipper” or “trigger” style can be used to gain initial entry. Rearrangement of the host cell actin cytoskeleton and polymerisation of F-actin at the site of the bacterial attachment is a common feature of these mechanisms (O Cróinín *et al*, 2012), while in phagocytic cells host medicated phagocytosisprocesses. Cytochalasin-D is a fungal toxin that binds tightly to the end of actin filaments thus acting as a potent inhibitor of actin polymerisation, and thus phagocytosis. In order to determine if actin rearrangement is important for the entry of *Photorhabdus* into host cells, THP-1 cells were treated with Cytochalasin-D prior to infection with either Kingscliff or PB68, using the methods consistent with the infection assays described above.

Inhibition of actin polymerisation by Cytochalasin-D caused a significant reduction in the abilities of both Kingscliff and PB68 bacteria to invade the cells (**Fig.6**). Perhaps not surprisingly, this differential effect was only seen when the bacteria were cultured at the temperature at which they had previously been shown to be most invasive at in the previous THP-1 invasion assays (**Fig 4**). For example, as described above, the Kingscliff strain can invade THP-1 cells but only when cultured at 37 °C prior to infection. When grown at 28 °C, the addition of Cytochalasin-D made no difference to any observed ability to “invade” cells. However, it should be noted that in these assays the recovery levels of the bacteria were approaching the lower range of sensitivity limit for the CFU counts, being only 1.6×10_1_. Strain PB68, on the other hand showed the converse of what was seen for the Kingscliff strain, with a significant reduction in invasion only at 28 °C. Nevertheless, while we did still see a trend for a drop in the number of recovered bacteria in the 37 °C grown samples for strain PB68, upon the addition of Cytochalasin-D, it was not significant.

**Fig. 6.**
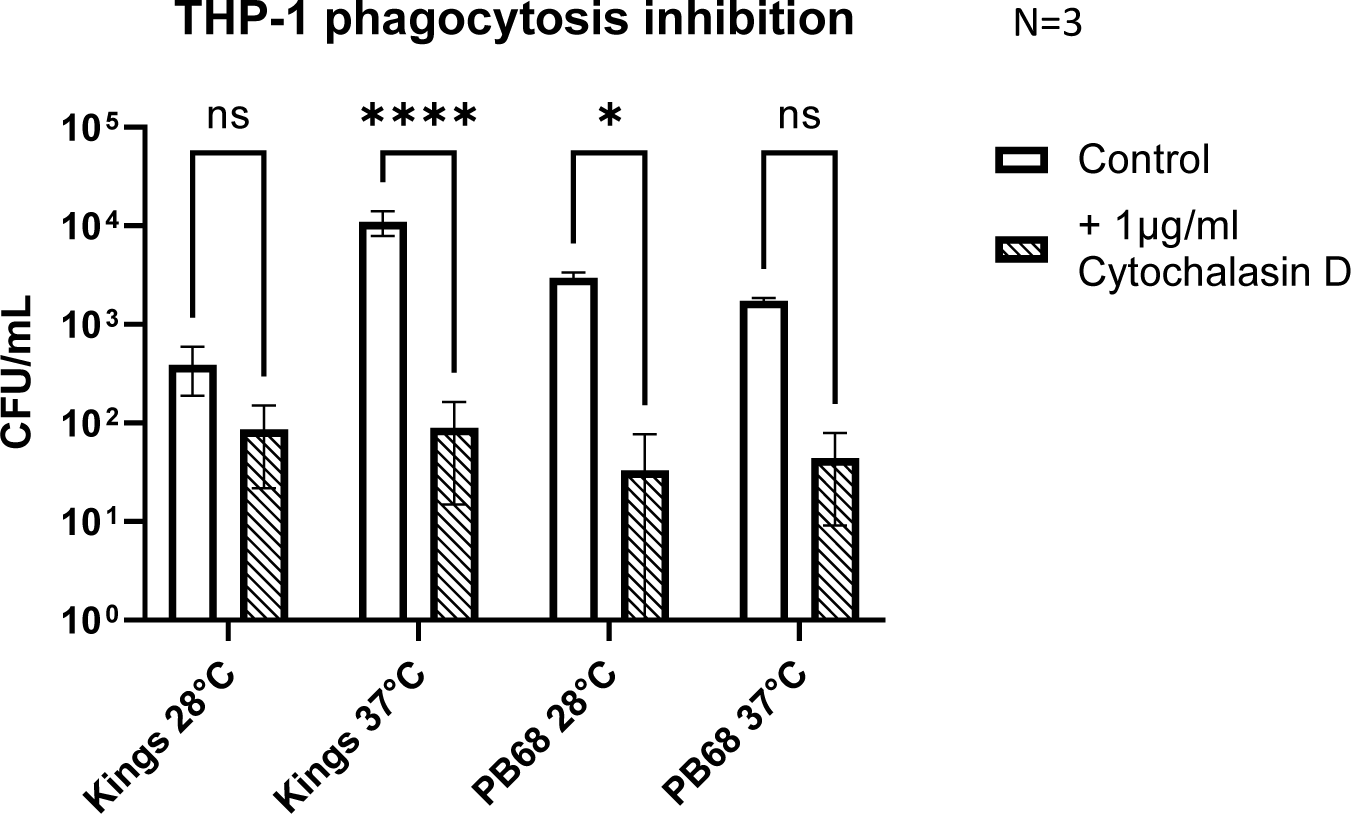
*Photorhabdus asymbiotica* subsp *Australis* strains, Kingscliff and PB68, with phagocytosis inhibited THP-1 cells. THP-1 cells were activated by PMA 48 hours prior to experiment. Bacteria were grown at either 28 °C or 37 °C prior to infection and allowed to interact for 2 hours. Prior to the infection assay phagocytosis inhibitor cytochalasin D was added to the THP-1 cells. (T-test, * = < 0.05, **** = < 0.0001, n = 3 biological replicates).

### Interactions of *Photorhabdus* with Peripheral Blood Mononuclear Cells (PBMCs)

The assays described above investigating *P. asymbiotica* interactions with mammalian cells were performed using immortalised cultured cells. Even in the case of cell lines such as the differentiated THP-1 macrophage like cells, this is not a perfect model of how the bacteria may behave when exposed to the more complicated and diverse human cellular immune system. A large part of the immune system response comes from the PBMCs which are round nucleated white blood cells found circulating in the blood. The PBMCs consist of; lymphocytes (T-cells, B-cells and NK-cells), monocytes and dendritic cells. These cell types represent both the early innate immune response, phagocytosis, antigen presentation activities by dendritic cells and tissue sequestered monocytes, and the late adaptive immune response, through the activation of T and B cells (and subsequent antibody production and immune memory). Thus, PBMCs taken from healthy human volunteers represent an excellent resource for studying the interactions of *P. asymbiotica* with a “natural” more complete cellular immune component model.

In order to determine which PBMC cell types, *P. asymbiotica* interacts with, either through adhesion or internalisation, constitutive GFP expressing strains of the; Australian (Kingscliff) and Thai (PB68) *P. asymbiotica* were used alongside a non-clinical *P. luminescens* strain (TT01-DJC), which provided a suitable negative control. Attempts were made to create a GFP expressing strain of the American *P. asymbiotica* (ATCC 43949) as well, but as with the HIT and JUN strains, it proved recalcitrant to transformation. The GFP expression strains were grown at either 28 °C or 37 °C, then allowed to interact with freshly harvested human PBMCs for 2 hours, before analysis using flow cytometry (**Fig.7, A**). To investigate if any of the observed interactions were either bacterial or PBMC activity dependant, control samples of these bacteria were killed prior to addition to the PBMCs. These controls allowed us to distinguish between passive phagocytic ingestion as opposed to active virulence processes. The flow cytometry allowed not only for the identification of the different PBMC cell types, through detection of differentiating surface markers (**Fig.7, B**), but also which were infected with *Photorhabdus* by identifying the GFP signal associated with each cell (**Fig.7, C**).

**Fig. 7.**
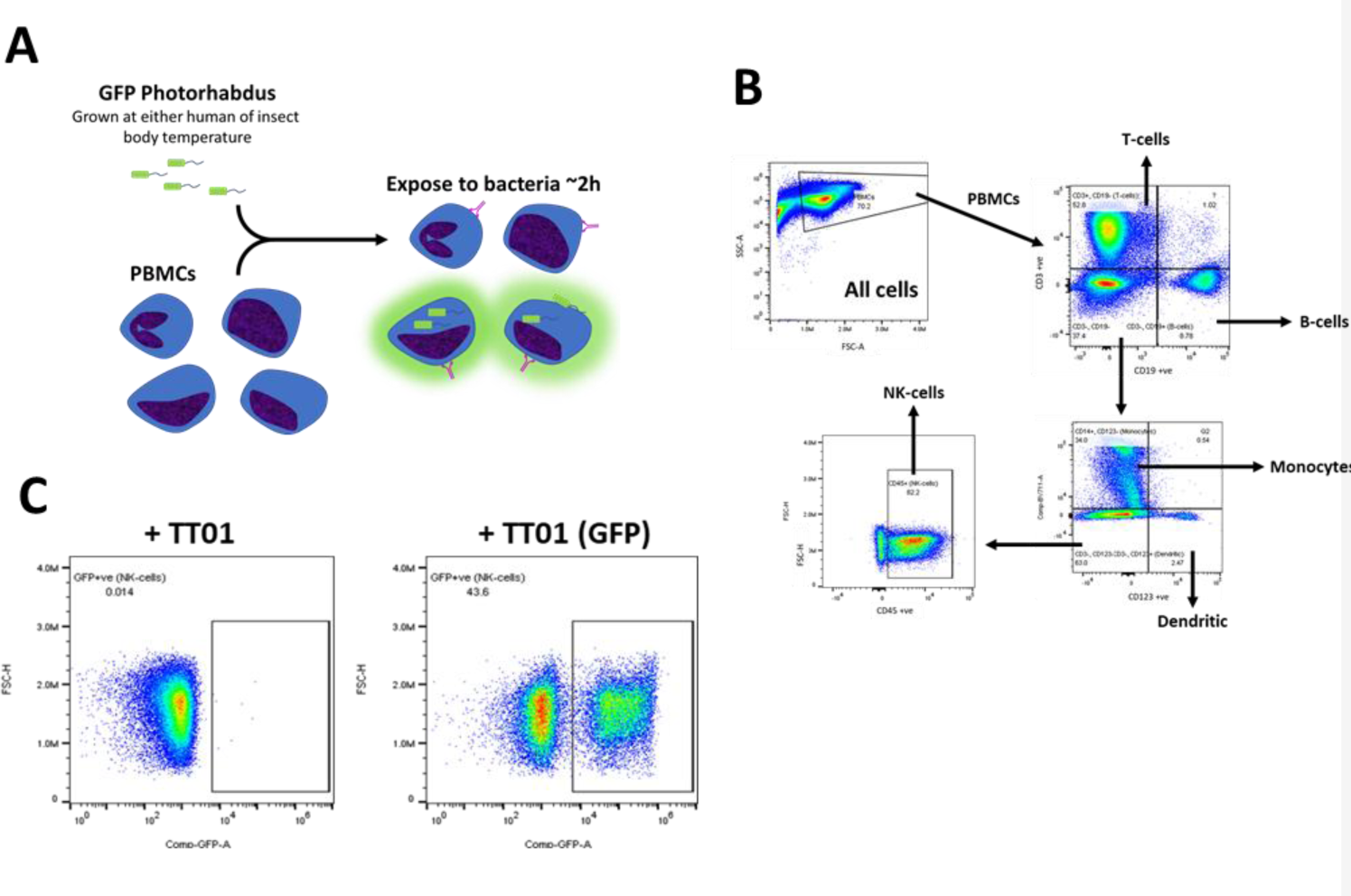
Flow cytometry strategy for identification of Photorhabdus infected PBMCs. **A-** Human derived PBMCs were exposed to various strains of *Photorhabdus* expressing GFP for 2 hours, before analysis by flow cytometry to calculate the percentage of PBMCs infected. In theory a GFP signal should be detected from PBMCs which have attached or internalised bacteria. The *Photorhabdus* strains were cultured first at either 28 °C (simulated ambient insect body temperature) or 37 °C (human core body temperature) prior to the infection. As controls *Photorhabdus* cultures of each strain were killed one hour before exposure to the PBMCs. **B-** Gating strategy for identifying the different PBMC cell types using flow cytometry. Antibody and fluorophore panel can be found in the methods section. **C-** Alongside detecting signals for each of the antibodies, once the PBMC types were gated out, GFP signal from any internal or attached bacteria was also detected. A distinct population of the cells (in this case monocytes) can be seen to have a GFP signal (+TT01_GFP), which is not seen when PBMCs are infected with non-GFP expressing bacteria (TT01). Using this a percentage of each PBMC cell type that exhibited a GFP signal, indicating infection with bacteria, could be calculated.

The findings from these experiments build upon our observations from the tissue culture invasion assays, indicating that the different *Photorhabdus* strains exhibit unique responses to the different PBMC cell types. Surprisingly, *P. asymbiotica* PB68 and *P. luminescens* TT01, showed very similar cell-type interaction profiles (**Fig.8 B, D**). Both had low levels of association with the majority of the PBMCs cell types, ∼0-30%, which dropped further for PB68 when grown at 38°C. There was also no significant change in the cell association profiles when the bacteria had been killed prior to exposure to the PBMCs, suggesting phagocytosis or passive cell surface binding is in operation. The only cell type that showed high levels of bacterial association were the dendritic cells, ∼70-90% at 28 °C. Importantly there was a significant drop for the pre-killed PB68 samples, though the number of GFP associated cells was still significantly higher than the other PBMCs cell types at this temperature, the drop was much larger for the TT01-DJC strain, dropping down to that of the other PBMC cell types.

**Fig. 8.**
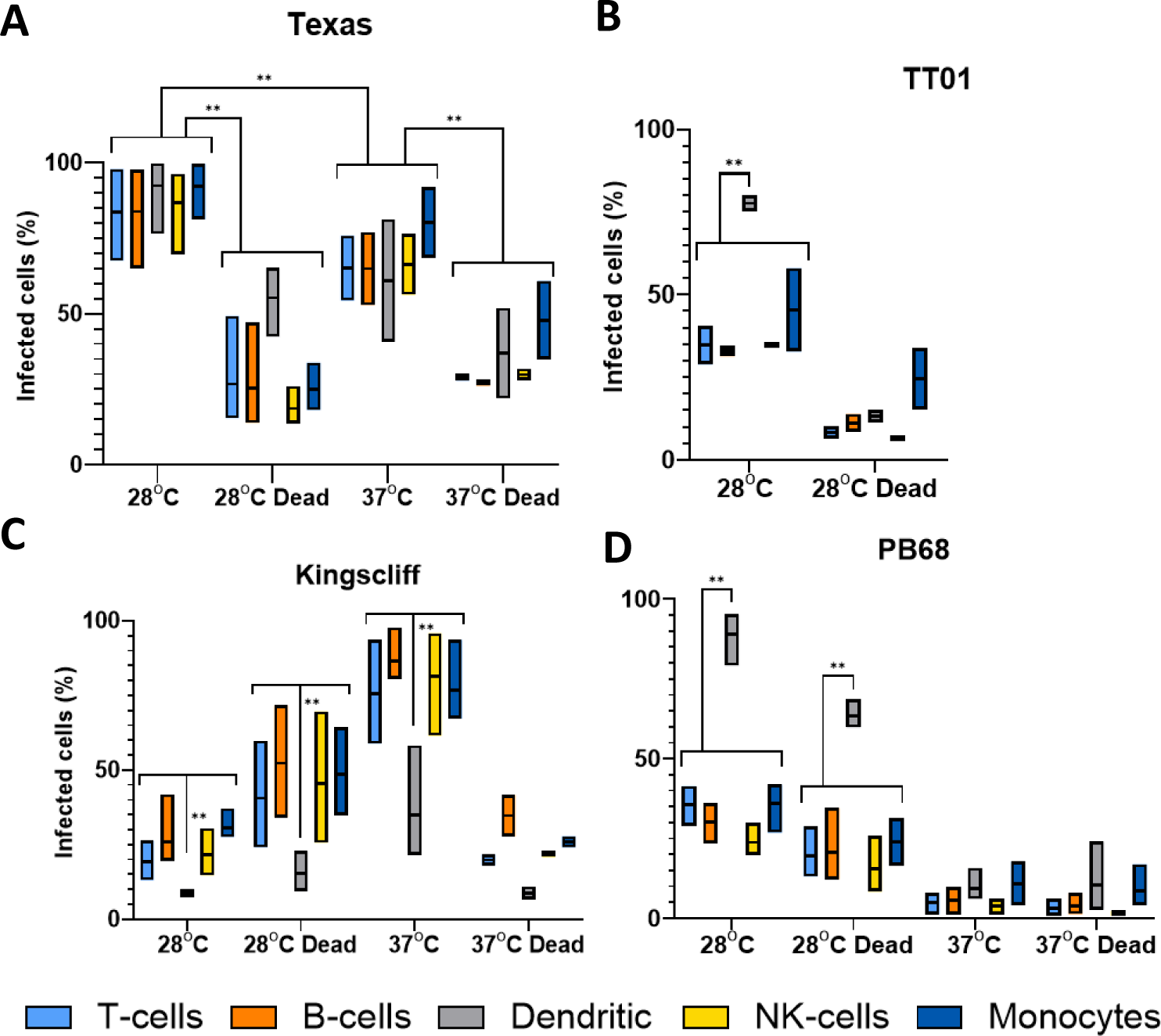
Flow cytometry analysis of varying infection rates of different stains of Photorhabdus. Human PBMCs were infected with different strains of GFP+ve *Photorhabdus* and analysed by flow cytometry as described in Fig.7, (3 experimental replicates for each sample). Four strains of *Photorhabdus* were tested; **A- Texas** the clinical *P. luminescens* isolate from a human neonate infection in the USA. **B-TT01-DJC** the *P. luminescens* lab strain originally isolated from a soil nematode. For this strain data could not be obtained for 38°C due to TT01_DJC’s inability to grow at this higher temperture. **C- Kingscliff** – a *P. asymbiotica* subsp. *Australis* isolate from a human infection in Australia, and **D- PB68** – a *P. asymbiotica* subsp. *Australis* nematode isolate from Thailand. (n = 2-4, 2-way anova, ** = < 0.01) Note: Only relevant P-values have been shown due to the number of statistical comparisons.

Unlike the other two strains tested, the Australian *P. asymbiotica* clinical strain (Kingscliff), showed very different behaviour in its associations with the various PBMC cell types. Bacteria grown at 28 °C generally showed low levels of cell association, ranging from 30-10% for most PBMC cell types. However, this increased significantly for Kingscliff when pre-cultured at 37 °C, reaching levels of 60-100%. When the 37 °C cultured Kingscliff strain was killed prior to exposure to the PBMCs, association rates dropped to a similar level seen for the live 28 °C grown bacteria, ∼40-10%. Strangely, the pre-killed bacteria grown at 28 °C, actually showed a slight increase over the live cells. Unlike the other *Photorhabdus* strains, at both cultured temperatures, even if the bacteria were killed prior to addition to the PBMCs, the dendritic cell association level was significantly lower than the other PBMCs. This contrasts with the other *Photorhabdus* strains where the dendritic cells always had association levels either the same, or higher than the other PBMC cell types.

Finally, the Texas strain again showed a different phenotype to all the other strains, both *P. asymbiotica* and *P. luminescens*. In this case all PBMC cell types showed high levels of bacterial association, with no significant distinction with dendritic cells unlike the other *Photorhabdus* strains. A higher proportion of the PBMCs had bacterial association when the Texas strain was grown at 28°C instead of 37°C, with all cell types almost reaching 100%. There was also a significant decrease when the bacterial were killed prior to incubation at both temperatures.

## Discussion

Here we have presented data obtained from both cultured and human derived ex vivo PBMC cells that details how prior bacterial growth temperature and host cell type strongly influence the behaviour of various strains of the *P. asymbiotica*. One of the first findings from these experiments is that the European HIT and JUN strains, which were confirmed by whole genome sequencing to be closely related genospecies to clinical isolates of *P. asymbiotica*, displayed no ability to survive within mammalian cells. Considering that they are relatively temperature intolerant, growth at 37 °C was unreliable and slow, and furthermore they have yet to be associated with a human infection, this perhaps should not have been unexpected. These observations, did indicate that they are not at all well adapted mammalian pathogens, unlike the known clinical isolates. These strains are likely either avoiding phagocytosis completely or are simply being killed by the THP-1 cells. Microscopic analysis of the interaction of these two strains with THP-1 cells suggested the former explanation for JUN and the latter for HIT which was seen to be internalised but not recovered during survival assays (Fig.4, 5). As JUN is not being internalised it suggests that it can inhibit the process such as through secreted effectors from the Type 3 Secretion System (T3SS). The T3SS deployment is a common mechanism that many Gram-negative pathogens use to manipulate lymphocytes and prevent phagocytic destruction (Santos et al, 2015). However, as JUN was not seen attaching to the THP-1 cells, this would not be possible for that strain. This suggests an alternative method JUN uses to evade internalisation. We speculate that HIT remains an exclusive insect pathogen, while JUN may have some limited potential as a mammalian pathogen. Nevertheless, these findings demonstrate that even bacteria isolated from similar geographic locations, for *P. asymbiotica* genospecies isolates, can show clear differences in their behaviour.

Previous research into the clinical isolates of *P. asymbiotica* showed that an American isolate was only weakly phagocytosed and could not invade Hela cells, unlike a representative Australian clinical isolate (Costa *et al*, 2009). Our data confirms these previous experiments, showing that our chosen American strain, ATCC 43949, shows only weak internalisation into either phagocytic or non-phagocytic cells, while the Australian Kingscliff strain showed high levels of internalisation (Fig.4, 7). The observation of internalisation in non-phagocytic cells for the Kingscliff strain suggests that cell invasion is an active virulence process. Conversely ATCC 43949’s low levels of internalisation in phagocytic cells suggests that it is instead actively evading of phagocytosis.

However, we expanded on these previous findings by also showing that these two strains differ in their response to growth temperature. Namely that ATCC 43949 behaviour is not affected by temperature, in contrast to Kingscliff which shows higher rates of internalisation when grown at human body temperature compared to the lower temperature more appropriate of an insect infection. These differences arise despite all incubations of the bacteria with the mammalian cells necessarily being performed at 37 °C. This confirms that prior growth at different temperatures leads to phenotypic adaptations that influence the outcome of the subsequent interactions for the Kingscliff strain. It is currently unknown exactly what changes in protein expression are responsible for these adaptations, but the most likely candidates would either be exotoxins, cell surface adhesins or capsular polysaccharides.

Interestingly, the genomes of the *P. asymbiotica* strains sequenced encode several homologs of *Yersinia* adhesion proteins, which could prove to be good subjects for further study. The selective advantage for temperature dependant cell tropisms for Kingscliff is likely related the need to conserve resources, as some of these pathogenicity factors are likely irrelevant for insect infections. Similar temperature dependant tropisms can be seen in many insect-human pathogens (such as *Yersinia pestis*), which in fact makes the American strain the more curious of the two strains (Konkel, 2000). It should also be noted that in the microscopic analysis of Kingscliff interactions, it is on occasion seen traversing a THP-1 cell membrane, suggesting the ability to enter or leave phagocytes though methods other than phagocytosis or cell lysis (**Fig. 5**). Taken together, these data suggest that the Kingscliff strain may have distinct regulons for both insect and mammalian infections.

While we were unfortunately not able to test how the American strain ATCC43949 interacts with human PBMCs, we were able to determine sound findings for the Kingscliff strain (Fig. 7). Once again there were large increases in bacterial association for nearly all PBMC types at the higher temperature, mirroring the behaviour seen in the cell line assays. However, the outlier were the dendritic cells, which at both temperatures showed significantly reduced bacterial association compared to the other PBMCs at either temperature. This is in contrast to the observations from the *P. luminescens* TT01-DJC strain and *P. asymbiotica* strain PB68, where dendritic cell bacterial association was generally higher than with other PBMC cell types. High levels of dendritic cell association would be expected as they are proficient professional phagocytes, therefore suggesting that Kingscliff may be actively evading the dendritic cells. Dendritic cells are also one of the key mediators of the early immune response, being responsible for presenting antigens, typically at the lymph-nodes, to activate T and B cell responses. Thus, by avoiding Dendritic cells Kingscliff may be delaying a rapid and lasting immune response, until it can establish a more robust infection. Surprisingly this difference was also observed with the pre-killed Kingscliff, indicating that the method of evasion is not an active one, and may be mediated by factors such as a bacterial capsule.

The fact that we have shown that there are large differences between the two human infective strains despite both showing similar symptoms such as distal secondary lesions, suggests that they have evolved different strategies to overcome the mammalian immune systems. This may not be surprising as the bacteria are geographically constrained by the need to inhabit either their soil nematode isolates or insect corpses, making migration from Australia to mainland North America unlikely.

One of the main symptoms of Photorhabdosis are secondary lesions formed distant from the primary site of infection, which is generally suggestive of a pathogen that is capable of dissemination through either the blood or lymphatic system, often by invading immune cells. A good example of this infection strategy is the plague bacterium *Yersinia pestis*, which can disseminate around the body by invading lymphatic system phagocytes. Our observations here suggest the Kingscliff strain may use a similar strategy, but as Kingscliff was seen to actively avoid dendritic cells it suggests it may preferentially invade non-phagocytic cell types. On the other hand, ATCC 43949 appeared to avoid internalisation, suggesting it may be using a strategy more similar to that of *Streptococcus pyogenes*, which instead attaches to the outside of immune cells as they travel to lymphatic nodes (Siggins et al, 2020).

One hypothesis as to why we see these differences in the American and Australian strains is that they may have different warm-blooded hosts. Humans are likely a dead-end host for *Photorhabdus*, the nematode vector being unable survive above 34 °C. However, it is possible that the nematode might be able to set up a successful infection alongside the *Photorhabdus* in a small warm-blooded animal such as a mouse if it is killed rapidly. If this is the case then varying warm blooded hosts in each region, may have led to the divergent evolution we see. Another possibility is even, that while the American strains infect mammals, the Australian strains instead infect ground dwelling birds. With this in mind it should be noted that only Kingscliff is able to readily grow up to a temperature of ∼42°C, which coincidentally is the average body temperature for most avian species.

Interestingly the recently identified first human pathogenic strain of *P. luminescens* (Texas strain), displayed different behaviour to both the American and Australian strains. Much like the ATCC 43949, growth temperature seemed to have no effect on invasion rates, this could be seen for both the THP-1 cells and PBMCs. However, in the THP-1 cells it exhibited a slightly higher rate of invasion than the American strain, although not to the same extent as the Australian strain. However, in the PBMC experiments the Texas strain showed a unique phenotype distinct from that of other strains where it was seen associating equally with all PBMC cell types at high levels.

This suggests that unlike the strategy used by the Kingscliff strain, which is to avoid dendritic cells, the Texas strain attacks all cell types, possibly attempting to overwhelm the immune system. This is reminiscent of how *Photorhabdus* acts in an insect infection, where the bacteria rapidly kill immune cells after establishing an infection, suggesting the Texas strain has adapted the more straight forward and aggressive approach used against the insect immune system to mammalian immune systems as well. While Kingscliff and possibly ATCC43949, have developed a distinct strategy for mammalian infection, which likely involves preventing early identification.

The final strain investigated was the Thai strain PB68, which up until this point had neither been studied nor shown to directly cause a human infection. In addition to the ability of the Thai strain to grow at 37 °C it also carries a homologue of the pPAU1 plasmid of ATCC43949, only found in the clinical isolates, which suggests it is indeed capable of mammalian infection. Backing up this hypothesis, unlike HIT and JUN, PB68 was able to survive challenge by the phagocytic THP-1 cells, and in fact showed higher rates of internalisation than the American strain, although not to the same level as Kingscliff. However, this could be brought into question by the observations of its interactions with the PBMCs, in which PB68 showed almost identical results to the non-human infective *P. luminescens* strain TT01-DJC. Both these bacteria had low rates of association in all cell types, apart from dendritic cells. Thus, unlike Kingscliff, these strains would likely get detected by the immune system early in the infection hampering their ability to successfully establish an infection in a mammalian host. Curiously a major difference between the Kingscliff and PB68 strains that further hints that it may not be a well-adapted mammalian pathogen relates to the observations that it reacted in the opposite way to Kingscliff in regard to activity upon previous growth temperature. While not causing a statistically significant increased growth rate at 28 °C, these conditions did lead to higher rates of invasion and association with both the cultured cells and the PBMCs. This, as is the case for the Kingscliff strain, showed that the bacteria are capable of responding to different host body temperatures. Although it is not known if this represents a strategy to “hide” from the immune system as we hypothesise for ATCC43949, or a down-regulation of pathogenicity genes which normally used for insect infections. A second difference that was noted in the behaviour of Kingscliff and PB68, was opposite preferences for invasion of cultured cell types. Where Kingscliff was by far the most successful in invading THP-1 and CHO cell lines, PB68 outperformed it by a large margin in the HEK cell line. Exactly what could be the cause of these differences, or the evolutionary implications is unknown. However, a likely candidate may be differing cell surface markers on these cell lines, which the bacteria are recognising.

In conclusion we have shown here that geographically distinct strains of *P. asymbiotica* not only differ in their ability to establish human infections, but also in their behaviour towards temperature and different host cell types. We also show that the novel human infective *P. luminescens* Texas strain can infect and survive inside human phagocytes. It also seems to aggressively colonise human PBMCs regardless of cell type or growth temperature, unlike the human infective *P. asymbiotica*. Considering this, it brings into question what molecular adaptations allow for a *Photorhabdus* species to become human infective. Knowing this will become more important over time as bio-pesticides, such as *Photorhabdus* and its symbiont nematode, see increasing popularity.

## Materials and Methodology

All materials and methods used during the studies found in this thesis have been collected into this section as to make for easier reading. Where appropriate in other chapters brief descriptions of methodology have been repeated to provide clarity and context.

**Table 1.**
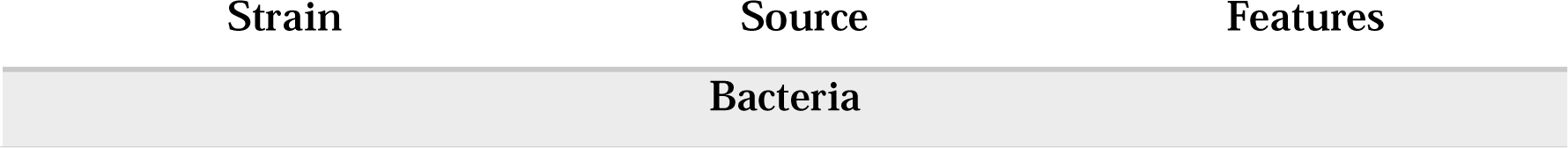
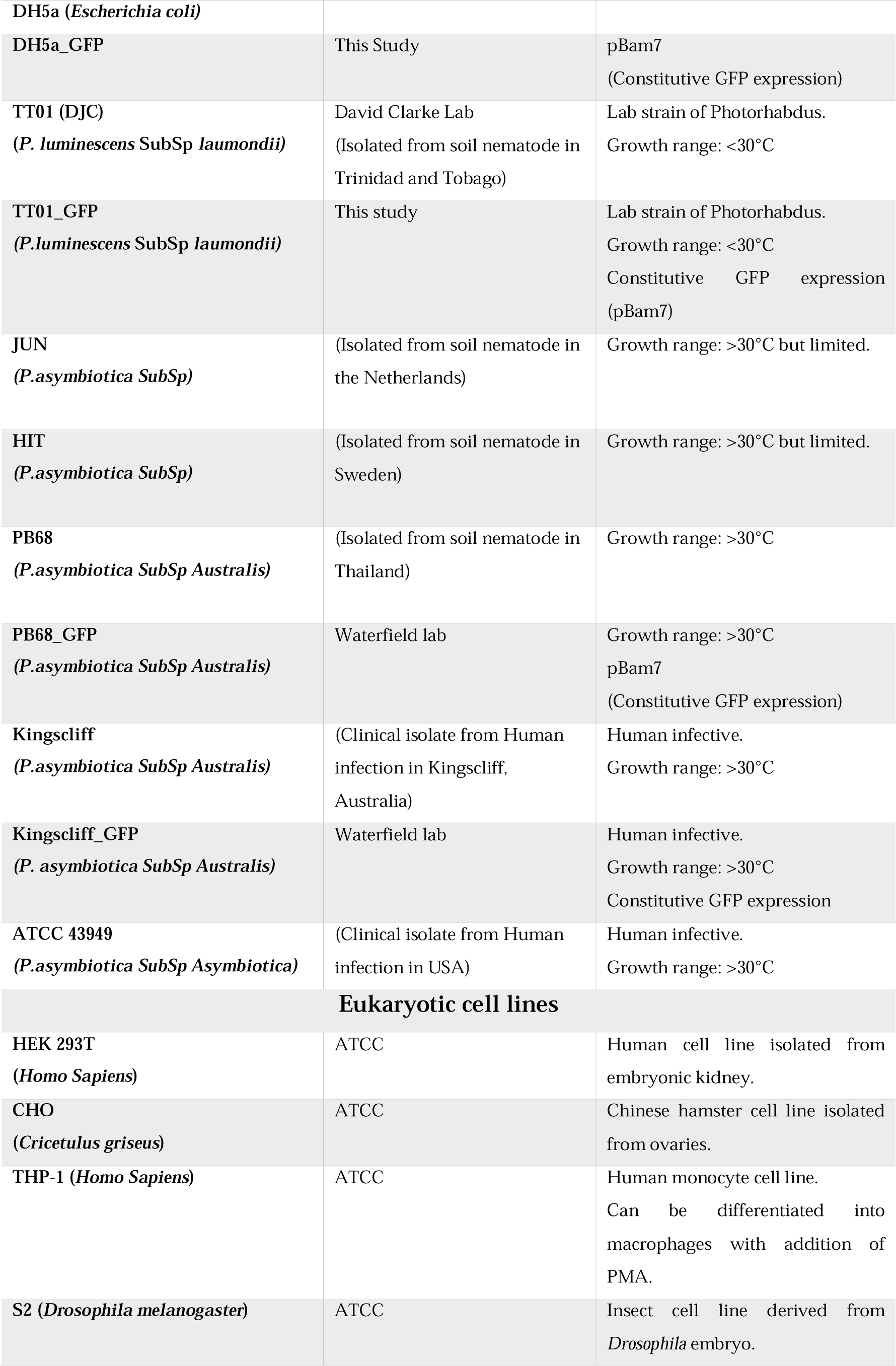
Bacterial and Cell lines used in this study.

### Bacterial protocols

#### Bacterial cultures

Routine growth of bacterial strains was carried out in standard lysogeny broth (LB), with shaking at ∼180rpm unless otherwise stated. *E. coli* cultures were grown at 37 °C, while *Photorhabdus* were grown at either 28 °C, or 37 °C, as required for the experiment. It should be noted that *P. luminescens* were normally grown 28 °C, due to having growth arrest above 34 °C. However, for some experiments a sub-culture of *P. luminescens* would be incubated at 37 °C, and while the bacteria remained viable, no further growth would occur. For growth on solid media, LB was supplemented with 1.5% agar, 0.1% sodium pyruvate and any relevant antibiotics; plates were incubated in the dark. *Photorhabdus* strains were confirmed to be sensitive to gentamycin prior to performing assays.

#### Antibiotics used

Various antibiotics were used during these studies for selection of transformed strains, the concentrations of which can be found in the table below (**Table 5.2**). It was observed that the

*P. asymbiotica* strains had a natural resistance to ampicillin, so this was avoided for use a selective agent in plasmids where possible.

**Table 2.** Common antibiotics and media supplements used during this study.

| USE | SUPPLEMENT | WORKING CONCENTRATION |
| --- | --- | --- |
| ANTIBIOTICS | Gentamycin | 10 µg/mL<sup>-1</sup> |
|  | Ampicillin | 100 µg/mL<sup>-1</sup> |
|  | Chloramphenicol | 25 µg/mL<sup>-1</sup> |
|  | Kanamycin | 25 µg/mL<sup>-1</sup> |
|  | PenStrep | 50 U/mL<sup>-1</sup> |
| PROMOTOR CONTROL | Arabinose | 0.2% (w/v) |
|  | Glucose | 0.2% (w/v) |

#### Transformations

All transformations of *E. coli* were done using the DH5a laboratory strain which was made chemically component to allow for introduction of plasmids through heat shock. Most of the transformations into *E. coli* were simply for the creation and replication of plasmids, due to its efficiency of plasmid uptake. *Photorhabdus* on the other hand does not seem to readily produce chemically competent cells. Thus, they were transformed using electroporation.

#### Electroporation of *Photorhabdus*

It has been found that unlike some cells, *Photorhabdus* loses competency when frozen, thus electrically competent cells must be created on the same day as the transformation. On the day of the transformation, 100ml of LB would be sub-cultured with 4ml of *Photorhabdus* grown overnight. This would be left to grow, as per normal *Photorhabdus* growth conditions, till ∼OD 0.2 (roughly 4 hours) then be placed on ice for 90 mins. The bacteria would then be pelleted at 4000xg, 10mins, 4°C then resuspended in 100ml of ice-cold SH buffer (5% [wt/vol] sucrose, 100 mM HEPES). The bacteria were the pelleted and resuspended in increasing smaller volumes of SH buffer; 50ml, 1.6ml, before being finally resuspended in 160µl of SH buffer. To pre-chilled 2mm Electroporation cuvettes, 40µl of the cells were added. While still on ice 4 µl of DNA was added to the cells and electroporation was carried out using the following parameters: 2.5 kV, 25 µF, and 200 Ω. After electroporation 1 mL of LB was quickly added to the bacteria and incubated in normal *Photorhabdus* growth conditions for 1h, after which they were plated onto LB plates with appropriate antibiotics.

### Eukaryotic cell protocols

#### Maintenance of cultured mammalian and insect cells

Maintenance of Mammalian cells was done at 37 °C with a humidified atmosphere of 10% CO_2_ with no shaking. HEK 293T and CHO cells were grown in DMEM media supplemented with 10% FBS, L-glutamine and non-essential amino acids. While THP-1 cells were grown in RPMI supplemented with 10% FBS, L-glutamine and non-essential amino acids. Cells were generally split at around 70-80% confluence.

#### Infection assays

Mammalian cells were seeded at 2×10_5_ onto sterilized glass coverslips in 24-well plates, in 500 µl of their respective cell media without antibiotics and left to adhere overnight at 37 °C with a humidified atmosphere of 10% CO_2._. In the case of THP-1 cells, 100nM of PMA was added after seeding 48 hours prior to the addition of bacteria to induce activation and adherence. Once the cells had fully adhered, bacteria grown to mid-log phase (OD 0.4-0.6) and resuspended in the cell media, were added to each well at a MOI of 1:50 (∼1×10_7_). Bacterial infection of cells was conducted over 2 hours, at 37 °C with a humidified atmosphere of 10% CO_2._ Afterwards 200 µg/ml of gentamicin was added to each well for 1h to kill any non-internalised bacteria. After incubation with the gentamycin, the media was subsequently removed and the cells were washed 3x with PBS, finally being resuspended in 100 µl of 1% Triton-X-100 (in PBS) for 10mins at room temperature, to lyse the cells and release any internalised bacteria. 900 µl of LB was then added and the cells were homogenised by pipetting. CFU counts were then done for each well as described above in the CFU/OD assay.

#### Inhibition of phagocytosis

THP-1 cells were seeded and activated as specified above for an infection assay, but 2 hours prior to addition of bacteria, 1 µg/mL of Cytochalasin D was added to the THP-1 cells. Afterwards the infection assay was carried out as normal.

#### Imaging of infected eukaryotic cells

Cells were grown and infected as with the infection assays above, but after the gentamycin incubation and washing steps, the cells were not lysed. Instead, the coverslips with the attached cells were transferred to new wells and fixed with 2% PFA (Paraformaldehyde) for 30 mins. After fixation, the cells were washed 3x with PBS and then had DNA stained using 300nM of DAPI for 5 mins. After staining, cells were again washed 3x with PBS and the coverslips were placed cells down onto a glass slide and sealed with clear nail varnish. Cells were then imaged using a Leica DMi8 fluorescence Microscope.

#### PBMC extraction and purification

Fresh whole human blood from healthy volunteers was diluted at a 1:1 ratio with PBS-EDTA and layered onto 12.5ml of ficoll medium in a 50 ml tube, making sure the blood and ficoll layer to not mix. The tube containing the ficoll/blood layers was spun at 400xg, room temperature for 30 mins, with acceleration and deceleration set at the minimum. This allowed the formation of a layer of PBMCs between the plasma and ficoll, which was carefully removed being sure to not take any of the other layers. The PBMCs were washed in PBS-EDTA and spun again at 300xg, room temperature, for 5 mins. The supernatant was removed and the pellet containing the PBMCs was resuspended in 20mL PBS-EDTA. The cells were then counted using a haemocytometer. PBMCs were then either used straight away or resuspended in RPMI at a dilution of around 20 million cells per mL for freezing. PBMCs for freezing after being resuspended had 2X freezing media added at 1:1 ratio and were aliquoted into cryotubes. The tubes were then either stored at −80 °C in Mr. Frosty freezing containers or in liquid nitrogen for long-term storage.

- 2X Freezing media
- – 20% DMSO in Foetal Bovine serum (FBS).

#### Flow cytometry of infected PBMCs

PBMCs were defrosted at 37 °C and resuspended in RPMI supplemented with 10% FBS and no antibiotics. Around 1×10_6_ PBMCs were seeded in wells for each condition/bacterial strain, including wells for the controls of no bacteria and unstained PBMCs. Bacteria, grown to mid log phase at either 28 °C or 37 °C, washed with PBS and resuspended in RPMI were added to the PBMCs at a MOI of 1:50 (cells:bacteria). For the killed bacterial assays, aliquots of the bacterial cultures were taken and killed with a mixture of 10ul/ml penstrep and 2ul/ml chloramphenicol for 1 hour prior to washing and addition to PBMCs. The bacteria and PBMCs were incubated together for 2 hours at 37 °C, after which 200 µg/mL of gentamycin was added for a further 1-hour incubation. After the gentamycin incubation the PBMCs were washed 3x in RPMI, then resuspended in 200ul of ice cold BSB and transferred to a 96-well V-bottom plate.

PBMCs were spun in the plate at 400xg for 5 mins, then resuspended in 50 µL of near-IR L/D stain (in PBS) and incubated in the dark on ice for 10 mins. PBMCs were washed in staining buffer and then resuspended in 25 µL of antibody cocktail (Table 4.1) and incubated in the dark on ice for 30 mins. PBMCs were then washed in 125µL BSB, then 150µL PBS and finally resuspended in 200 µL Staining buffer.

Stained PBMC samples were analysed by flow cytometry using a Cytek Aurora, with compensations having been previously generated for each antibody-fluorescence channel. The GFP signal from the bacteria was detected using the FITC channel on the instrument, with compensations for this channel being generated using PBMCs stained with a GFP conjugated antibody. This antibody was not included in the final experimental panel.

**Table 2.** Antibodies used for identification of PBMCs in flow cytometry. Cell marker represents the cell surface protein target of the specific antibody, and the florescent dye is the conjugated fluorophore.

| CELL MARKER | FLORESCENT DYE |
| --- | --- |
| CD45 | Pacific Orange |
| CD3 | AF-700 |
| HLA-DR | BV-786 |
| CD56 | PE-Cy7 |
| CD19 | APC |
| CD14 | BV711 |
| CD16 | BV605 |
| CD11C | BV421 |
| CD123 | PE |
| L/D | Near-IR |

### Molecular Techniques

All DNA, plasmids and primers were stored at −20 °C and kept on ice when in use.

**Table 3.**
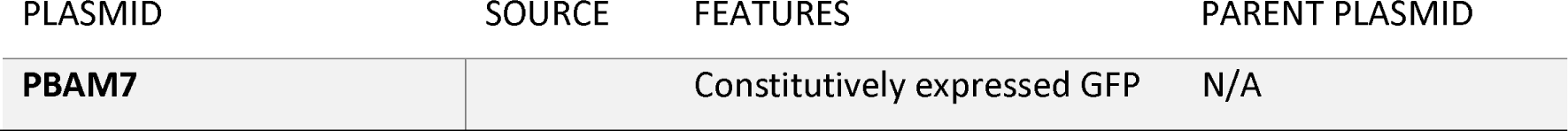
Plasmids used in this study.

#### Purification of Plasmids

Plasmids were purified from bacterial strains using the Qiagen Miniprep Spin Kit as per the manufacturer’s instructions. 5 mL overnight cultures were used, and the final elution was conducted in 2x 20 µL washes using molecular grade water.

#### Taq and colony PCR

When sequence reliability was not a concern such as during colony PCR, or for very small amplicons Taq polymerase was used. PCR parameters and relative volumes of reagents were done per manufacturers recommendations. For rapid screening of possible transformants, a small amount of individual bacterial colonies were taken from plates and boiled in 50 µL water at 95°C for 5 mins. This was then used as the template for the colony PCR.

#### Agarose gel electrophoresis

Quantification and identification of DNA fragment sizes was done using Agarose gel electrophoresis. 1% gels (w/v) were added to 1x Tris-Acetate-EDTA (TAE) buffer and the mixture was microwaved until the agarose had melted. The solution was allowed to cool slightly then SYBER-safe gel stain was added at a dilution of 1:10000. The agarose was poured into a mould and a left to set for around 20 mins after which the gel was loaded with the sample, alongside the GeneRuler 1kb Plus Ladder from Themo-Fisher, and the gel was run at 110 V for around 40-45 mins. Visualization of the gel was done using the Bio-Rad ChemiDoc MP Imaging System.

#### Primers

All primers used in these experiments were ordered from IDT.

**Table 4.**
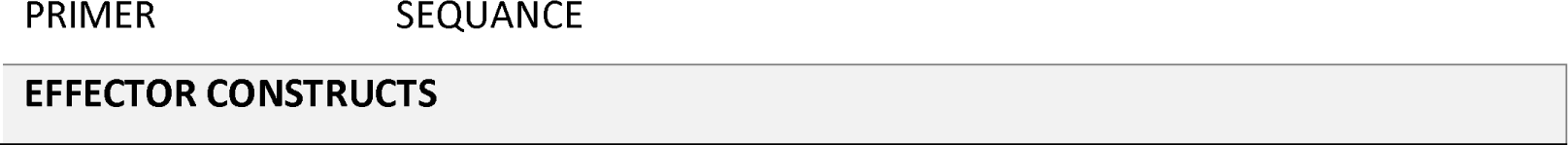
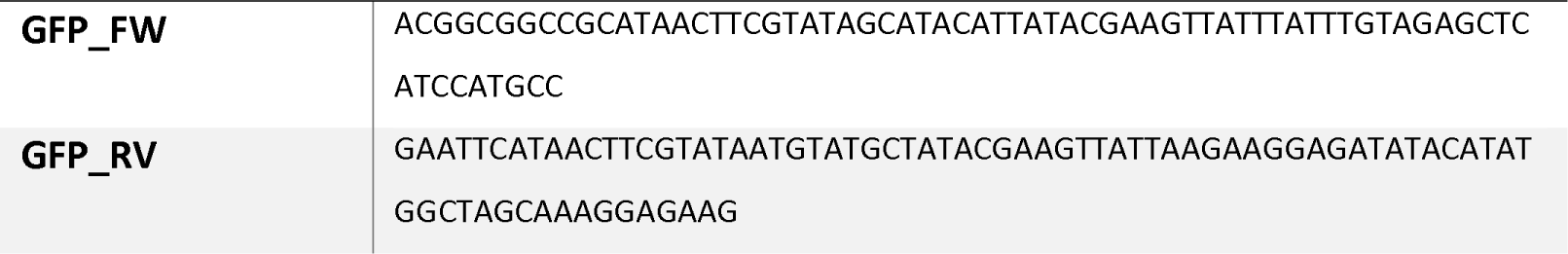
Primers used in this study.

#### Statistical tests

All statistical tests along with P-values used are found in the figure legends of the appropriate figure. Unless otherwise stated all tests were done in prism using standard settings.

